# An essential role for dNTP homeostasis following CDK-induced replication stress

**DOI:** 10.1101/395715

**Authors:** Chen-Chun Pai, Kuo-Feng Hsu, Samuel. C. Durley, Andrea Keszthelyi, Stephen E. Kearsey, Charalampos Rallis, Lisa K. Folkes, Rachel Deegan, Sarah E. Wilkins, Sophia X. Pfister, Nagore De León, Christopher J. Schofield, Jürg Bähler, Antony M. Carr, Timothy C. Humphrey

**Author notes:** Present Address: Kuo-Feng Hsu, Department of Surgery, Tri-Service General Hospital, National Defense Medical Centre, Taipei, Taiwan.

## Abstract

Replication stress is a common feature of cancer cells, and thus a potentially important therapeutic target. Here we show that CDK-induced replication stress is synthetic lethal with mutations disrupting dNTP homeostasis in fission yeast. Wee1 inactivation leads to increased dNTP demand and replication stress through CDK-induced firing of dormant replication origins. Subsequent dNTP depletion leads to inefficient DNA replication, Mus81-dependent DNA damage, and to genome instability. Cells respond to this replication stress by increasing dNTP supply through Set2-dependent MBF-induced expression of Cdc22, the catalytic subunit of ribonucleotide reductase (RNR). Disrupting dNTP synthesis following Wee1 inactivation, through abrogating Set2-dependent H3K36 tri-methylation or DNA integrity checkpoint inactivation results in critically low dNTP levels, replication collapse and cell death, which can be rescued by increasing dNTP levels. These findings support a ‘dNTP supply and demand’ model in which maintaining dNTP homeostasis is essential to prevent replication catastrophe in response to CDK-induced replication stress.

## Introduction

Replication stress, in which DNA replication forks stall, is a source of genome instability and a common feature of cancer cells (Gaillard, Garcia-Muse, & Aguilera, 2015). The ability to target such a hallmark of cancer cells is of significant therapeutic interest. Replication stress can result from multiple events including physical blockage of replication fork progression, deregulation of the replication initiation or elongation complexes, or through deoxyribonucleotide triphosphate (dNTP) depletion (Dobbelstein & Sorensen, 2015; Zeman & Cimprich, 2014). Cells respond to such events by triggering checkpoint dependent responses to facilitate DNA replication restart (Mazouzi, Velimezi, & Loizou, 2014). In humans ATR and CHK1 are the primary kinases responsible for replication checkpoint activity, while in fission yeast the Rad3 and Cds1 kinases play a predominant role, with Cds1 being redundant with Chk1 in this response (Boddy, Furnari, Mondesert, & Russell, 1998; Feijoo et al., 2001; Flynn & Zou, 2011; Howard D. Lindsay et al., 1998). Unresponsive stalled forks can be subject to endonucleolytic cleavage by Mus81-Eme1, generating a DNA end, which is targeted for homologous recombination (HR) (Hanada et al., 2007; Roseaulin et al., 2008).

In fission yeast, dNTP synthesis is induced in response to replication stress and DNA damage by at least two distinct mechanisms (Guarino, Salguero, & Kearsey, 2014). Checkpoint activation promotes Ddb1-Cul4^Cdt2^-dependent degradation of Spd1, an inhibitor of ribonucleotide reductase (RNR), thereby promoting dNTP synthesis (Holmberg et al., 2005; Liu et al., 2005; Liu et al., 2003). In addition, checkpoint-dependent activation of the MluI Cell Cycle Box (MCB) binding factor (MBF) complex promotes transcription of genes encoding one or more MCB domains within their promoter regions, including *cdc22*^*+*^, the catalytic subunit of RNR, thereby promoting dNTP synthesis (Dutta et al., 2008).

The chromatin state plays an important role in modulating transcriptional responses. Set2 is a histone methyltransferase required for histone H3 lysine 36 (H3K36) mono, di- and trimethylation in yeast (Morris et al., 2005). Various functions have been ascribed to H3K36 methylation, including DNA repair (Pai et al., 2014) and checkpoint signalling (Jha & Strahl, 2014). Further, we recently described a role for Set2 in promoting dNTP synthesis in response to DNA damage and replication stress through promoting MBF-dependent transcriptional expression of *cdc22*^*+*^. Loss of Set2 leads to reduced Cdc22 expression, resulting in reduced dNTP levels and consequent replication stress (Pai et al., 2017). Such roles for Set2 in maintaining genome stability help explain the tumour suppressor function of the human orthologue, SETD2.

Replication stress can also arise as a result of elevated CDK activity, and Cyclin E and Cyclin A are frequently overexpressed in cancers (Hwang & Clurman, 2005; Yam, Fung,&Poon, 2002). Wee1 is a negative regulator of cell cycle progression where it phosphorylates and inactivates Cdc2/CDK1 kinase, thereby preventing entry into mitosis (Russell & Nurse, 1987). Inactivation of Wee1 upregulates CDK activity and promotes G2-M progression. In addition to regulating entry into mitosis, studies in mammalian cells have found that WEE1 kinase inhibition can lead to dNTP depletion through increased firing of replication origins resulting from deregulated CDK activity (Beck et al., 2012).

Synthetic lethality provides an opportunity to specifically target cancer cells (Chan & Giaccia, 2011). In this respect, previous studies using fission yeast identified checkpoint mutants (r*ad1Δ, rad3Δ, rad9Δ, rad17Δ, hus1Δ)* that are synthetic lethal with Wee1 inactivation using a temperature sensitive allele of Wee1, *wee1-50* (al-Khodairy & Carr, 1992; Enoch, Carr,&Nurse, 1992). These *wee1-50* checkpoint deficient double mutants manifest a strong ‘cut’ (<u>c</u>ell <u>u</u>ntimely <u>t</u>orn) phenotype in which the genetic material is mis-segregated into daughter cells, consistent with cell death arising from mitotic catastrophe (Enoch et al., 1992). Indeed, inhibitors to human WEE1 have been developed with the aim of promoting mitotic catastrophe in G1-S checkpoint deficient p53 mutant cancer cells (Hirai et al., 2009). As the synthetic lethal relationship between Wee1 inactivation and loss of Chk1 is conserved in mammalian cells (Chila et al., 2015), and because inhibitors to human WEE1, ATR and CHK1 have been developed with the aim of targeting cancer cells (Dobbelstein & Sorensen, 2015; Sørensen & Syljuåsen, 2012), understanding the mechanism by which their inactivation leads to cell death is of clinical significance.

In this study, we define an evolutionarily conserved role for Wee1 in preventing replication stress through suppressing CDK-induced replication origin firing, dNTP depletion and Mus81-dependent DNA damage. Further, we show that following Wee1 inactivation, Set2-dependent histone H3K36 trimethylation and the DNA integrity checkpoint perform an essential role in maintaining dNTP homeostasis, thus preventing replication catastrophe. These findings provide new insights into the consequences of Wee1 inactivation and its therapeutic exploitation.

## Results

### Wee1 is required for efficient S-phase progression by limiting origin firing

We investigated the possible role of Wee1 in regulating S-phase progression. Nitrogen starvation was used to synchronize *wee1-50* cells in G1 phase and following re-feeding, cell cycle progression was monitored by flow cytometry. In wild-type cells, an increasing proportion of cells with a 2C DNA content was observed at 3 hours following re-feeding; by 5 hours, the entire population was 2C, indicating successful DNA replication (Fig. 1a, **WT**). In contrast, in *wee1-50* cells, at 3 hours after re-feeding the population exhibited a 1C peak, and even 6 hours following re-feeding there was a proportion of *wee1-50* cells with a 1C peak, indicating a delay in S-phase progression (Fig. 1a, ***wee1-50***).

**Figure 1.**
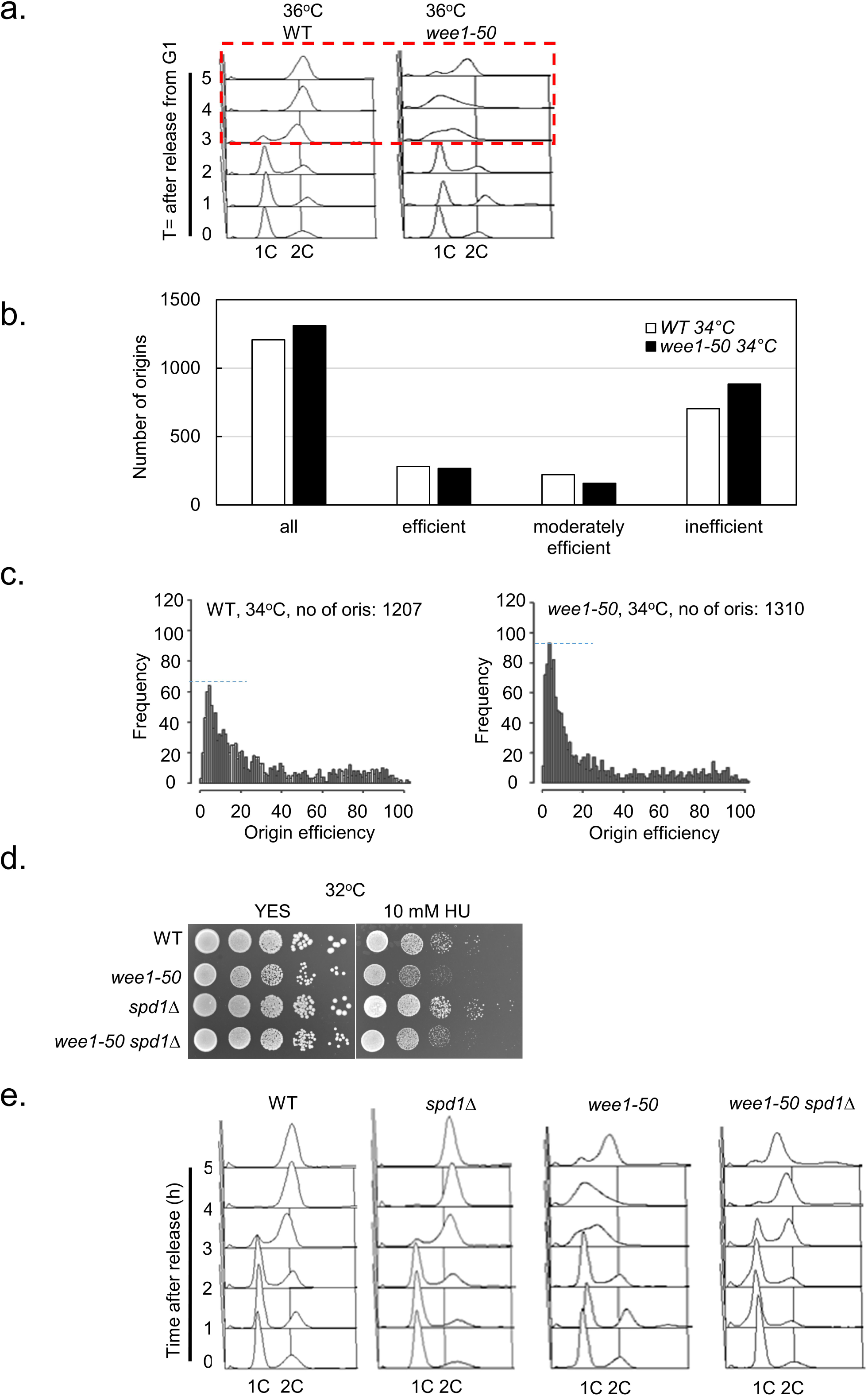
Wee1 suppresses dormant origin firing and dNTP depletion. (**a**) Wee1 is required for efficient DNA replication. Log phase wild-type or *wee1-50* cells were blocked in G1 phase by nitrogen starvation in EMM-N for 16h at 25°C. Cells were released from the G1 block by re-suspending in EMM+N at 36°C. Samples were collected at the indicated time points for FACS analysis. The red dashed line box indicates the delayed S phase progression in *wee1-50* cells. (**b**) Wee1 suppresses inefficient origin firing. The genome-wide plot of origin usage in *wee1-50* cells in comparison with wild-type cells at 34°C. Origin efficiencies were calculated from Pu-seq data [25]. The sequencing experiment was performed once and therefore it is not possible to perform a statistical analysis. (**c**) The quantification of the frequency of origin usage (efficiency) in asynchronous wild-type and *wee1-50* cells at 34°C. (**d**) Spd1 depletion suppresses the sensitivity of *wee1-50* cells to HU. WT and *wee1-50* cells were serially diluted and spotted onto YES plates containing 10mM HU and incubated at 32°C for 2-3 days. (**e**) Deletion of *spd1*^*+*^ promotes S-phase progression in *wee1-50* cells. Wild-type, *spd1Δ, wee1-50* and *spd1Δ wee1-50* cells were arrested in G1 by nitrogen starvation, released and samples taken at time points indicated and subjected to FACS analysis.

To test whether Wee1 inactivation in fission yeast causes increased origin firing, we employed a polymerase usage sequence (Pu-seq) technique to map genome-wide origin usage as previously described (Daigaku et al., 2015). In wild-type cells, we identified 1,207 initiation sites at 34°C (threshold 20 percentile, 99.9 percentile of all origins regarded as 100% efficient) including efficient (>50% usage per cell cycle), moderately efficient (25-50%), and inefficient origins (<25%) (Fig. 1b). In the *wee1-50* background, we mapped 1,310 origins at 36°C (Fig. 1b). Interestingly, analysis of the distribution of origin usage in *wee1-50* cells revealed the trend that an increased number of inefficient origins (dormant origins) were used compared to wild-type cells (Fig. 1b). There are a greater proportion of inefficient origins and less efficient origins in *wee1-50* cells compared to wild type (Fig. 1c). Together, this data suggests that Wee1 inactivation causes an increase in the number of DNA replication initiation sites utilized.

We tested whether the increased origin firing in *wee1-50* might lead to elevated dNTP demand, thus leading to replication stress. A spot assay showed that *wee1-50* cells were sensitive to HU at the semi-restrictive temperature (Fig. 1d and **Supplementary Fig. 1a)**. Deleting RNR inhibitor *spd1*^+^ in a *wee1-50* background suppressed the sensitivity of *wee1-50* cells on HU (Fig. 1d and **Supplementary Fig. 1a)** and suppressed the delayed DNA replication of *wee1-50* cells at 36°C, consistent with Wee1 inactivation impacting on dNTP levels (Fig. 1e). Consistent with this, we showed that the dATP/ATP level in *wee1-50* is significantly lower than wild type (**Supplementary Fig. 1b)**. These findings suggest that inactivation of Wee1 causes dNTP pool depletion by increased origin firing leading to replication stress.

### Wee1 inactivation causes DNA damage accumulation and genome instability

We next tested whether disrupting Wee1 could lead to DNA damage associated with replication stress. We monitored DNA damage-induced Rad52 foci in a *wee1-50* mutant. Wee1 inactivation resulted in significantly elevated levels of Rad52 foci compared to wild-type (Fig. 2a and 2b). Earlier work has demonstrated that increased CDK activity promotes Mus81-Eme1 endonuclease activity (Dehe et al., 2013; Dominguez-Kelly et al., 2011). Indeed, deletion of *mus81*^*+*^ resulted in significantly reduced levels of Rad52-GFP DNA damage foci in a *wee1-50* background (*p* value <0.05) (Fig. 2c and 2d). Thus, Wee1 inactivation leads to elevated levels of Mus81-dependent DNA damage.

**Figure 2.**
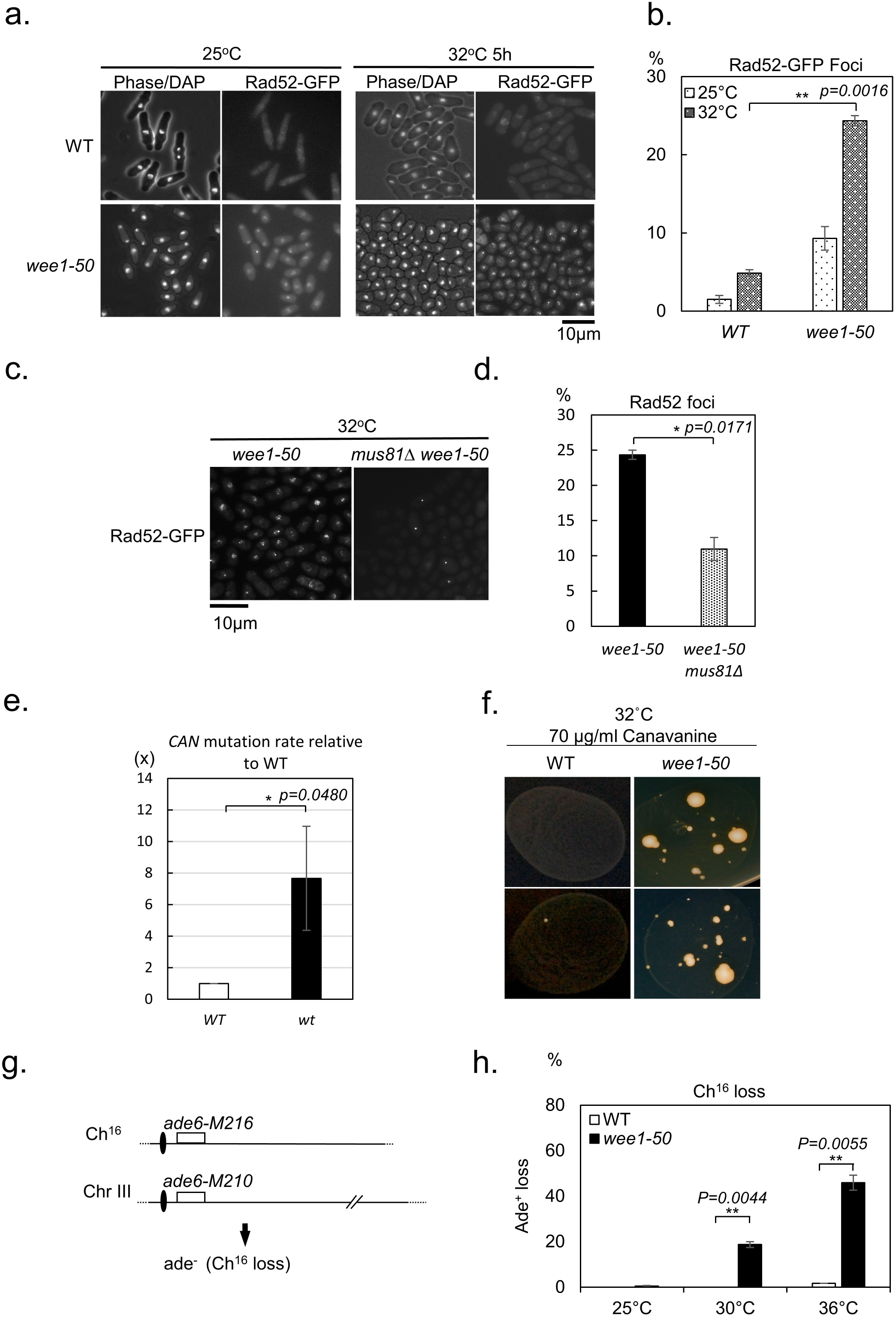
Wee1 inactivation causes DNA damage, increases mutation rates and leads to Ch^16^ loss. (**a**) Examination of Rad52-GFP foci in WT or *wee1-50* cells at 25°C or 32°C. Cells were grown to log phase at the permissive temperature before transferring to the semi-permissive temperature for 5h. Samples were fixed directly in methanol/acetone and examined by florescence microscopy. (**b)** The percentage of cells containing Rad52-GFP foci in the indicated strains is shown. A total of >100 cells were counted in each experimental group in two independent experiments (** *t* test, p<0.01). (**c**) A similar experiment was carried out as described in (**a**), except a *wee1-50 mus81Δ rad52-GFP* stain was used. (**d**) Quantification analysis of *wee1-50 mus81Δ* cells with Rad52-GFP foci compared to *wee1-50* cells. A total of >100 cells were counted in each experimental group in two independent experiments (* *t* test, p<0.05). (**e**) *wee1-50* cells exhibit elevated mutation rates to Can^r^ compared to wild-type cells. Cell cultures incubated on canavanine plates at 32°C for 10 days produced Can^r^ mutant colonies. Colony data were collected from 36 independent cultures. The mutation rates for WT and *wee1-50* strains were calculated using the MSS statistical method. The mutation rates and error bars are shown (averages of n ≥ 2 experiments, * *t* test, p<0.05). (**f**) The images presented are representative of experiments performed in (**e**) at least three times. (**g**) Schematic of the Ch^16^ strain. Ch^16^, ChIII, centromeric regions (ovals), complementary heteroalleles (*ade6-M216* and *ade6-M210*). (**h**) Elevated Ch^16^ loss rates associated with Wee1 inactivation. Wild-type or *wee1-50* cells containing the mini-chromosome are *ade*^*+*^. Cells were plated on YES or adenine-limiting plates and the percentage of Ch^16^ loss events per division was determined (n >500 cells for each data point, ** *t* test, p<0.01). The data presented are from at least two independent biological repeats.

Studies in budding yeast have shown that dNTP imbalance can cause mutagenesis and induce genome instability (Kumar et al., 2011). Therefore, we tested whether Wee1 inactivation associated with DNA damage or dNTP deregulation induces mutagenesis. We used resistance to canavanine (Fraser, Neill, & Davey, 2003; Kaur, Fraser, Freyer, Davey, & Doetsch, 1999) to determine the mutation rate in wild-type and *wee1-50* backgrounds. Inactivation of Wee1 showed significantly higher mutation rates (*p* value <0.05) compared to wild type (Fig. 2e and 2f).

It is known that either increasing or decreasing origin efficiency increases the loss of minichromosome Ch^16^ due to effects on replication fork stability (Patel et al., 2008). Consistent with this, *wee1-50* cells displayed high rates of minichromosome Ch^16^ loss at the semi-restrictive (30°C) or restrictive temperature (36°C) compared to wild-type cells (Fig. 2g and 2h). Together, these results suggest that Wee1 is essential for maintaining genome stability through suppressing replication stress, which leads to DNA damage, mutagenesis, and replication fork collapse.

### Loss of Set2 methyltransferase activity is synthetic lethal with wee1-50

Given that the histone H3K36 methyltransferase Set2 is required for DSB repair (Pai et al., 2014) and MBF-dependent transcription in response to DNA damage (Pai et al., 2017), we tested the possibility that the double mutant *set2****Δ*** *wee1-50* would be sick due to the accumulated DNA damage caused by Wee1 inactivation (Fig. 2a and 2b). Consistent with this, the *set2****Δ*** *wee1-50* double mutant was synthetic lethal when grown at the restrictive temperature (36°C) (Fig. 3a). To determine whether this synthetic lethality was dependent on the histone methyltransferase activity of Set2, *wee1-50* was crossed with a *set2* mutant (*set2-R255G*) in which the methyltransferase activity was abolished (Pai et al., 2014). The *set2-R255G wee1-50* double mutant was not viable at the restrictive temperature of 36°C (Fig. 3b), indicating that the methyltransferase activity of Set2 is required for viability in the absence of Wee1 kinase. Accordingly, *wee1-50* was also synthetic lethal with a *H3K36R* (Fig. 3c). Taken together, these results imply loss of Set2-dependent H3K36 methylation is synthetic lethal with Wee1 inactivation.

**Figure 3.**
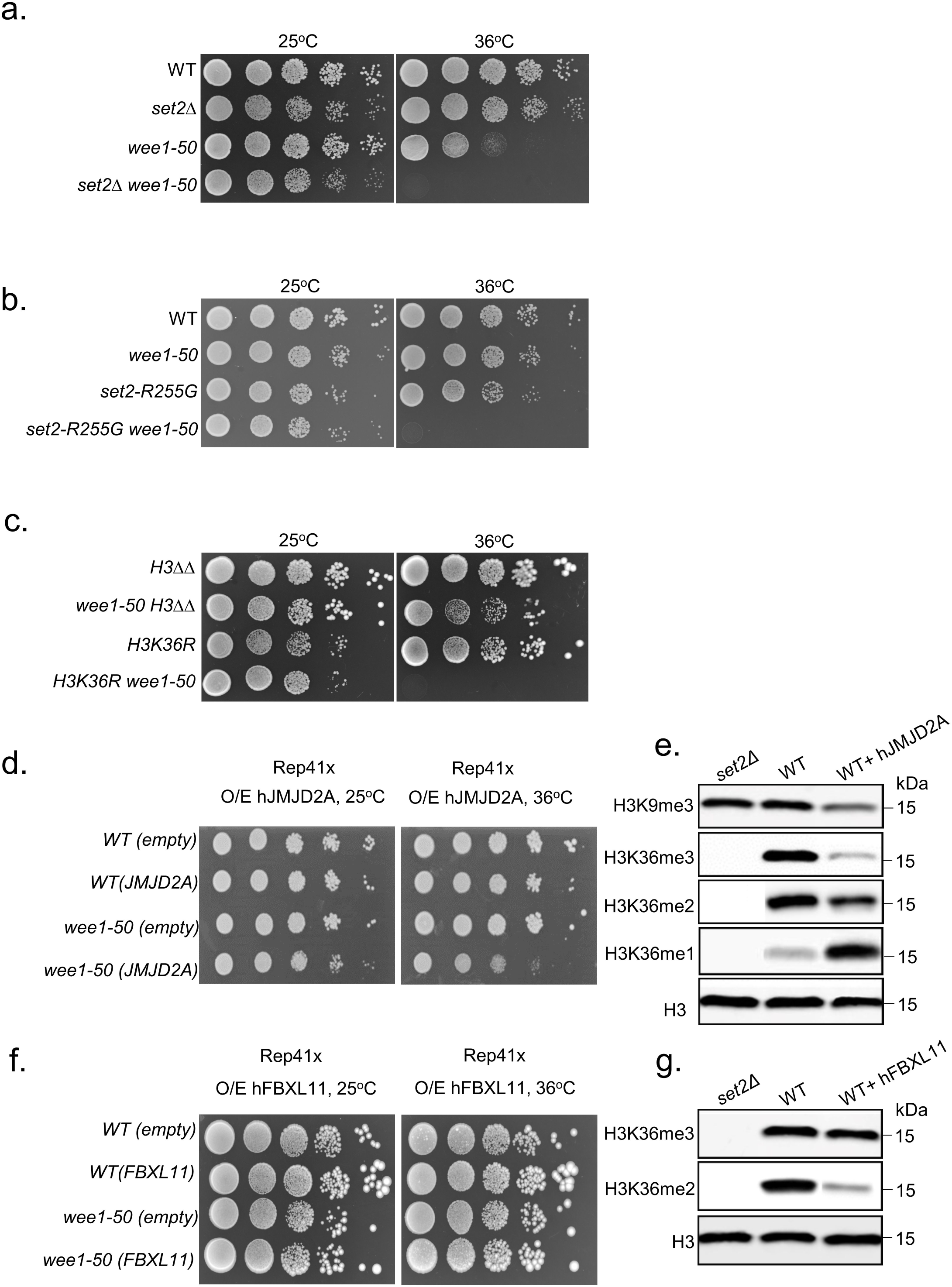
Loss of Set2-dependent H3K36 methylation is synthetic lethal with *wee1-50*. (**a**) WT, *set2Δ, wee1-50* and *set2Δ wee1-50* cells were serially diluted and spotted onto YES plates and incubated at indicated temperatures for 2-3 days. (**b**) WT, *set2-R255G, wee1-50* and *set2-R255G wee1-50* cells were serially diluted and spotted onto YES plates and incubated at indicated temperatures for 2-3 days. (**c**) H3*ΔΔ, wee1-50, H3K36R, H3K36R wee1-50* cells were serially diluted and spotted onto YES plates and incubated at indicated temperatures for 2-3 days. (**d**) Serial dilutions of wild-type cells (WT) expression empty vector *pREP41x* or *pREP41x-JMJD2A*, and *wee1-50* mutants expressing empty vector *pREP41x* or *pREP41x-JMJD2A*. Transformants were serially diluted and spotted onto EMM minus Leucine in the absence of thiamine at 25°C or 36°C. (**e**) Western blotting analysis of H3K9me3, H3K36me3, H3K36me2 and H3K36me1 in wild type (WT) containing *pREP41x* or *pREP41x-JMJD2A* and *set2Δ* cells. H3 is shown as a loading control. (**f**) Serial dilutions of wild-type cells (WT) expression empty vector *pREP41x* or *pREP41x-FBXL11*, and *wee1-50* mutants expressing empty vector *pREP41x* or *pREP41x-FBXL11*. Transformants were serially diluted and spotted onto EMM minus Leucine in the absence of thiamine at 25°C or 36°C. (**g**) Western blotting analysis of H3K36me3 and H3K36me2 in wild type (WT) containing *pREP41x* or *pREP41x-FBXL11* and *set2Δ* cells. H3 is shown as a loading control.

### Loss of H3K36 tri-methylation is synthetic lethal with wee1-50

In contrast to SETD2, the human homologue, Set2 in *S. pombe*, is responsible for all three forms of H3K36 methylation (H3K36me1, 2 or 3) and thus its loss cannot be used to distinguish between methylation states (Morris et al., 2005). We therefore investigated the consequences of expressing hJMJD2A/KDM4A (here termed hJMJD2A), the human demethylase that catalyzes H3K36me3/me2 to H3K36me2/me1, under the control of the thiamine repressible (*nmt*) promoters (**Supplementary Fig. 2a** and **2c**), on a plasmid in wild-type or *wee1-50* cells at the permissive or restrictive temperatures (Hillringhaus et al., 2011; Klose et al., 2006; Shin & Janknecht, 2007; Whetstine et al., 2006). Moderate overexpression of hJMJD2A was synthetic sick with *wee1-50* at 36°C (Fig. 3d, and **Supplementary Fig. 2e**). Consistent with previous studies, expression of human JMJD2A resulted in reduction of H3K36me3 and H3K36me2 levels **(Fig. 3e)**. In addition to H3K36me3 loss, expressing hJMJD2A also resulted in reduced levels of H3K9me3 (Fig. 3e and **Supplementary Fig. 2c**). However, as we did not observe synthetic lethality between deletion of *clr4*^*+*^, encoding the H3K9 methyltransferase, and *wee1-50* (**Supplementary Fig. 3**), this indicates that H3K9me3 loss is not required for cell viability in the absence of Wee1.

To distinguish between loss of H3K36me3 and H3K36me2, we expressed wild-type human H3K36me2-specific demethylase JHDM1A/KDM2A/FBXL11 (hFBXL11) in wild-type or *wee1-50* cells (**Supplementary Fig. 2b**, **2d** and **2f**) (Tsukada et al., 2006). Accordingly, we found that expression of hFBXL11 in fission yeast resulted in significant decrease in H3K36me2 but did not affect H3K36me3 levels (Fig. 3g), indicating that hFBXL11 preferentially demethylates H3K36me2 *in vivo*. However, expression of hFBXL11 did not induce a significant viability loss in *wee1-50* cells at 36°C (Fig. 3f), and expression of hJMJD2A or hFBXL11 did not sensitize wild-type or *wee1-50* cells at the permissive temperature (Fig. 3d and **3f**). Collectively, these findings provide strong evidence that the histone mark H3K36me3 is required for viability in the absence of Wee1.

### set2Δ synthetic lethality with wee1-50 can be suppressed by Cdc2 inactivation

We next explored whether Wee1 inactivation leads to synthetic lethality with *set2Δ* through elevated CDK activity or through a CDK independent function. To test this, we investigated whether we could suppress the synthetic lethality by inhibiting CDK activity. We crossed the analogue-sensitive *cdc2* mutant (*cdc2-as*) (Dischinger, Krapp, Xie, Paulson, & Simanis, 2008) with *set2****Δ*** *wee1-50* to create a *cdc2-as set2****Δ*** *wee1-50* triple mutant. Instead of using the ATP analogue molecule (1-NM-PP1) to inactivate Cdc2 activity, we found that the *cdc2-as* mutant exhibited modest temperature sensitivity. As shown in **Supplementary Fig. 4a**, loss of CDK activity suppressed the growth defect of *set2****Δ*** *wee1-50* mutants. Further, the triple mutant showed a plating efficiency of 87.5 ± 1.5% as compared to *set2****Δ*** *wee1-50,* which was only 0.3 ± 0.3% (**Supplementary Fig. 4b**) while *set2****Δ*** and *wee1-50* single mutants exhibited more than 90% plating efficiency. Collectively, these results indicate elevated CDK activity resulting from Wee1 inactivation leads to synthetic lethality in a *set2Δ wee1-50 background.*

**Figure 4.**
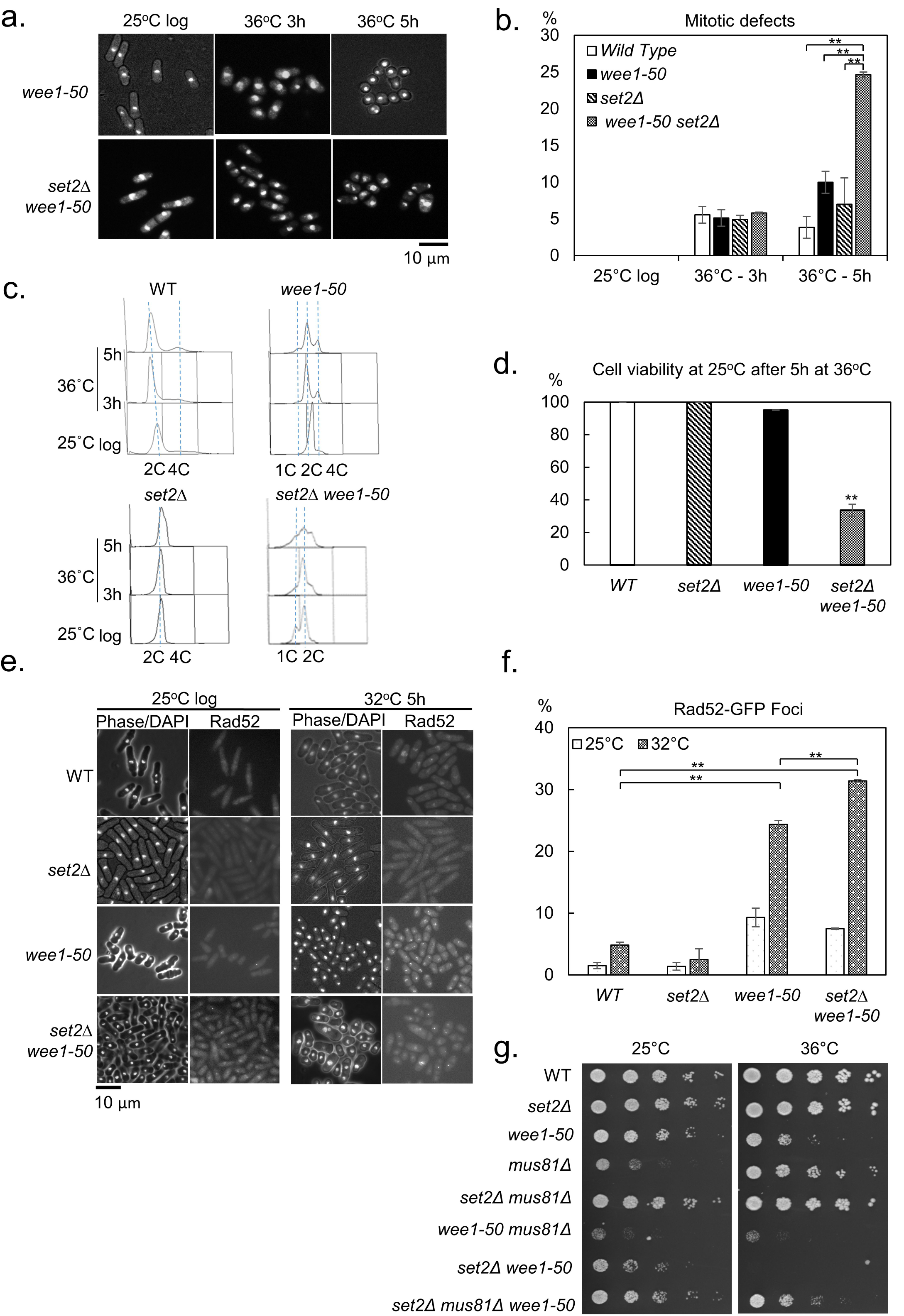
*set2Δ* synthetic lethality with *wee1-50* results from replication catastrophe. (**a**) Inactivation of Wee1 in *set2Δ* cells results in premature entry into mitosis. *wee1-50* or *set2Δ wee1-50* cells were grown to log phase at permissive temperature (25°C), then incubated at 36°C to inactivate Wee1. Samples were fixed with 70% ethanol at indicated times. The fixed cells were stained with DAPI and examined by microscopy analysis. (**b**) Quantitative analysis of cells in (**a**). Asterisks represent significant differences (n ≥ 2 experiments for each genotype, n >200 cells for each data point; ** *t* test, p<0.01). The data presented are from at least two independent biological repeats. (**c**) A wild-type, *wee1-50, set2Δ* or *set2Δ wee1-50* strain was grown to log phase at the permissive temperature 25°C then transferred to 36°C for the times shown. At the indicated times, cells were processed for FACS analysis. (**d**) WT, *set2Δ, wee1-50* and *set2Δ wee1-50* cells from the 5h time point in (**c**) were collected, plated on the YES medium and incubated at 25°C for 3-4 days for viability analysis. (n ≥ 2 experiments for each genotype, n>500 cells for each data point; ** *t* test p<0.01, t test between WT and *set2Δ wee1-50* cells: p-value = 0.0031). (**e**) Examination of Rad52-GFP foci in WT, *set2Δ, wee1-50* or *set2Δ wee1-50* cells at 25°C or 32°C. Cells were grown to log phase at the permissive temperature before transferring to the semi-permissive temperature for 5h. Samples were fixed directly in methanol/acetone and examined by florescence microscopy. (**f)** The percentage of cells containing Rad52-GFP foci in the indicated strains is shown. Asterisks represent significant differences (** *t* test p<0.01; averages of n ≥ 2 experiments, n ≥100 cells for each data point). (**g)** Deletion of *mus81*^*+*^ partially suppresses the synthetic lethality of *set2Δ wee1-50* cells at 36°C.

### set2Δ synthetic lethality with wee1-50 results from replication catastrophe

Previous studies using fission yeast found that a number of checkpoint mutants (r*ad1Δ, rad3Δ, rad9Δ, rad17Δ, hus1Δ)* were synthetic lethal with *wee1-50* at the restrictive temperature (al-Khodairy & Carr, 1992; Enoch et al., 1992). These double mutants exhibited a ‘cut’ (<u>c</u>ell <u>u</u>ntimely torn) phenotype suggesting that cell death arose through mitotic catastrophe (Enoch et al., 1992). Thus, we suspected that the synthetic lethality seen in *set2****Δ*** *wee1-50* cells might be also due to premature entry into mitosis. We found that *set2****Δ*** *wee1-50* cells were ‘wee’, and 25.6% of *set2****Δ*** *wee1-50* cells exhibited a ‘cut’ phenotype at 36°C after 5h incubation (Fig. 4a and 4b). This level of cutting in *set2****Δ*** *wee1-50* cells was significantly higher than *wee1-50* cells (10%) (*p* value <0.05) (Fig. 4a and 4b). Surprisingly, *set2Δ wee1-50* cells showed a striking S-phase delay even at the permissive temperature (Fig. 4c), suggesting inactivation of Wee1 causes more extreme DNA replication defects in *set2Δ* cells. Flow cytometry analysis showed that *set2Δ wee1-50* cells accumulated in S-phase following a shift to 36°C for 3-5 h (Fig. 4c). This result suggests that the observed cell death might be substantially due to a permanent replication stalling in the *set2Δ wee1-50* double mutant rather than through mitotic catastrophe (Fig. 4a and 4b). Nevertheless, 72 ±5 % of *set2****Δ*** *wee1-50* cells exhibited a ‘cut’ phenotype following a shift to 36°C for 24h (**Supplementary Fig. 5a** and **5b**), suggesting that the majority of *set2****Δ*** *wee1-50* cells eventually undergo mitotic catastrophe after long-term replication stalling.

To investigate whether the replication arrest was the cause of synthetic lethality in *set2Δ wee1-50* cells, double mutants were incubated at the restrictive temperature of 36°C for 5 h and the cell viability was examined by returning them to the permissive temperature of 25°C. The results showed that 66.3% of *set2Δ wee1-50* cells lost viability after shifting to the restrictive temperature for 5 h (Fig. 4d), in which majority of double mutants had arrested during DNA replication but only 26% of the double mutants underwent mitotic catastrophe (Fig. 4b), suggesting that most *set2Δ wee1-50* cells were dying in S-phase.

Further, we found that *set2Δ wee1-50* cells exhibited elevated levels of DNA damage compared to wild-type cells (Fig. 4e and **4f**), indicative of replication stress-induced DNA damage accumulation. Accordingly, *mus81*^*+*^ deletion resulted in partial suppression of the *set2Δ wee1-50* synthetic lethality at 36°C (Fig. 4g). In this respect, as *mus81*^*+*^ deletion had little effect on *set2Δ* viability, Mus81-dependent cleavage in the double mutant is likely to have arisen from Wee1 inactivation alone. In contrast, we also observed that Mus81 is required for *wee1-50* viability at 36°C. Together, these data suggest that Wee1 inactivation in a *set2Δ* background leads to elevated levels of replication fork collapse and to Mus81-dependent DNA cleavage.

### set2Δ wee1-50 replication catastrophe results from nucleotide depletion

Our data indicate that Wee1 inactivation leads to nucleotide depletion. Further, we have independently identified a role for Set2 in dNTP synthesis. We showed that dNTP levels were lower in *set2Δ* cells compared to wild type (Pai et al., 2017). We therefore tested the possibility that the *set2Δ wee1-50* synthetic lethality during S-phase was due to severe nucleotide depletion. Consistent with this, we found *set2Δ wee1-50* cells to be acutely sensitive to low levels of HU (Fig. 5a and 5b). These cells exhibited elongated phenotypes with HU treatment, suggesting that the lethality of the double mutant was not due to the compromised checkpoint (**Supplementary Fig. 6**). Instead, the lethality was more likely to be due to dNTP starvation. Consistent with this, we found that Cdc22 levels (the catalytic subunit of RNR) were reduced in response to replication stress induced in a *set2Δ wee1-50* double mutant compared to a *wee1-50* mutant (Fig. 5c). In accordance with this observation, dNTP levels are also significantly lower in *set2Δ wee1-50* in comparison to *wee1-50* cells under replication stress at 36°C (*p* value <0.005) (Fig. 5d). Further, deleting *spd1*^*+*^, encoding a negative regulator of RNR (Hakansson, Dahl, Chilkova, Domkin, & Thelander, 2006; Liu et al., 2003) robustly suppressed the synthetic lethality of the *set2Δ wee1-50* double mutant at 36°C (Fig. 5e), indicating elevated dNTPs can suppress *set2Δ wee1-50* synthetic lethality. Consistent with this observation, we found that dNTP levels are higher in the *spd1Δ set2Δ wee1-50* triple mutant compared to *set2Δ wee1-50* double mutants (Fig. 5f). Together, these findings indicate that the *set2Δ wee1-50* synthetic lethality resulted from dNTP depletion, to below a critical level.

**Figure 5.**
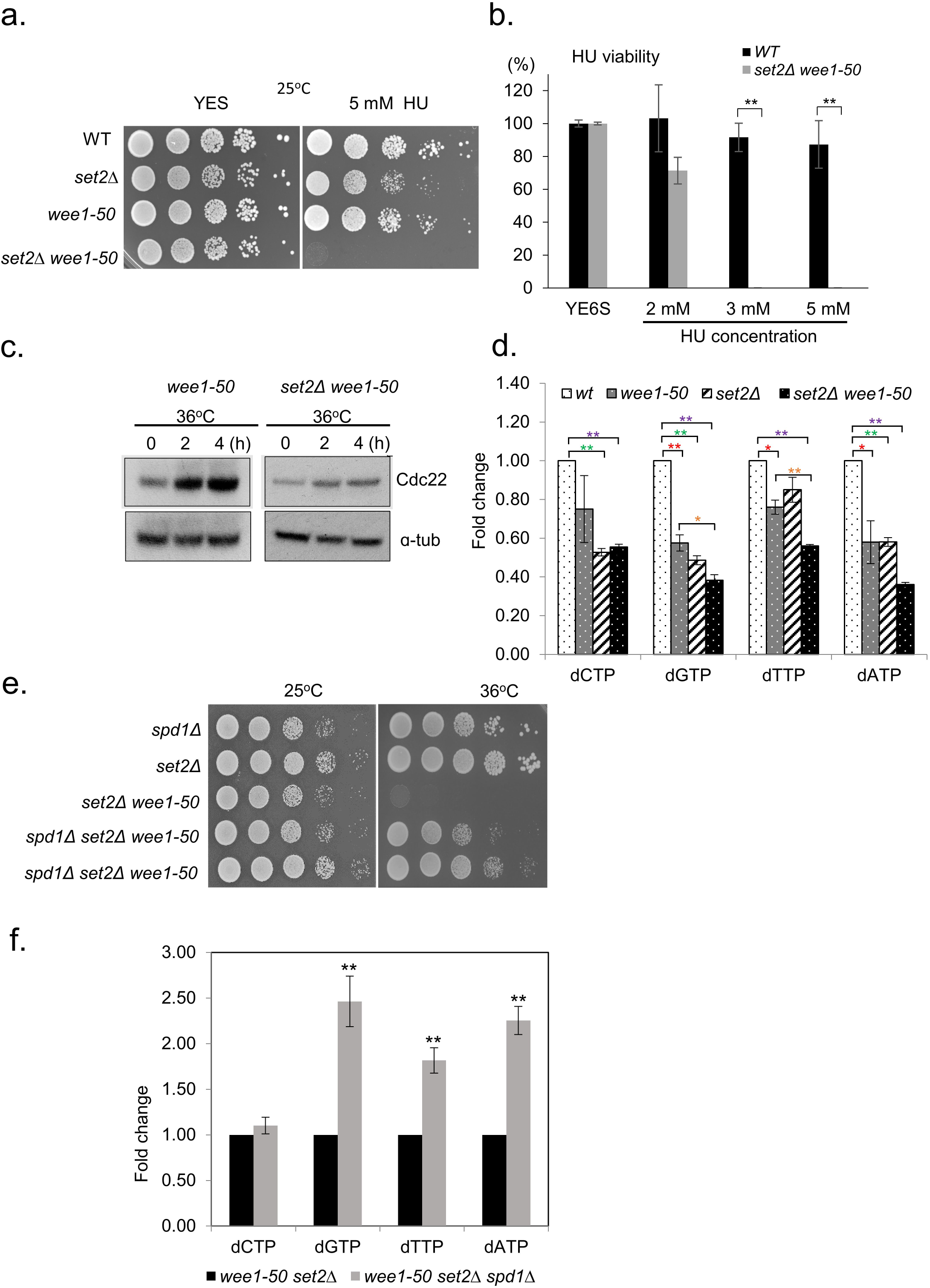
synthetic lethality of *set2Δ wee1-50* results from dNTP depletion. (**a**) *set2Δ wee1-50* cells are sensitive to low levels of HU. WT, *set2Δ, wee1-50* and *set2Δ wee1-50* cells were serially diluted and spotted onto YES plates containing 5mM HU and incubated at the permissive temperature (25°C) for 3-4 days. (**b**) Quantification of the viability of wild-type and *set2Δ wee1-50* cells on YES plates at 25°C containing different concentrations of HU as indicated (** *t* test p<0.01; averages of n ≥ 2 experiments, n ≥500 cells for each data point). (**c**) The protein levels of Cdc22 were examined in *wee1-50* and *set2Δ wee1-50* cells following incubation at 36°C for 4h. Samples of cells were taken at the indicated time points and cell extracts were made using TCA method. Cdc22 was detected using an antibody against the CFP tag. α-tubulin is shown as a loading control. (**d**) dNTP levels were measured in wt, *wee1-50, set2Δ* and *set2Δ wee1-50* strains. Cells were grown to log phase at 25°C followed by 5h incubation at 36°C. Samples of cells were collected and re-suspended in 10% TCA for subsequent HPLC analysis following neutralisation. Means ± standard errors of three experiments are shown. Stars denote statistical significance (* *t* test p<0.05, ** *t* test p<0.01). (**e**) *spd1Δ* suppresses the synthetic lethality of *set2Δ wee1-50*. Strains were serially diluted and spotted onto YES plates and incubated at indicated temperatures for 2-3 days. (**f**) dNTP levels were measured in *set2Δ wee1-50* and *spd1Δ set2Δ wee1-50* strains. Cells were grown to log phase at 25°C followed by a 5h incubation at 36°C. Samples of cells were collected and re-suspended in 10% TCA for subsequent HPLC analysis following neutralisation. The mean ± standard error for three experiments are shown. Asterisk(s) represents significant differences (** *t* test p<0.01, *t* test between *set2Δ wee1-50* and *spd1Δ set2Δ wee1-50* strains; p-values: dCTP=0.3288, dGTP=0.0065, dTTP=0.0042, dATP=0.0011). The data presented are from three independent biological repeats.

### set2Δ wee1-50 nucleotide depletion is caused by down-regulation of the transcription of MBF-dependent genes and increased origin firing

We have shown that Set2 controls dNTP synthesis through regulation of MBF transcription activity (Pai et al., 2017). As part of that study we found that deletion of MBF transcriptional repressor Yox1 suppressed the prolonged S-phase in *set2Δ* cells (Pai et al., 2017). Thus, we tested whether deletion of Yox1 could suppress the lethality of the *set2Δ wee1-50* double mutant and found that the triple mutant exhibited an increase in viability (Fig. 6a), indicating elevated MBF transcription activity can suppress *set2Δ wee1-50* synthetic lethality, presumably due to increased dNTP pools. Consistently, deletion of MBF transcriptional repressor Nrm1 also supressed the synthetic lethality of *set2Δ wee1-50* cells (Fig. 6b). Further, we tested whether Set2 also affected mRNA levels of MBF-dependent genes in a *wee1-50* background. To do this, the *set2Δ wee1-50* double and *wee1-50* single mutants were grown at the restrictive temperature of 36°C for 5h and global levels of gene expression were compared using microarrays. This analysis revealed that transcription of the MBF-dependent genes *tos4*^*+*^, *cdt1*^*+*^ and *mik1*^*+*^ was reduced following *set2*^*+*^ deletion in a *wee1-50* background, while *act1*^*+*^ which is not MBF-induced, was not (Fig. 6c). Together, these findings support a role for Set2 in facilitating MBF transcription in response to DNA damage or replication stress resulting from Wee1 inactivation.

**Figure 6.**
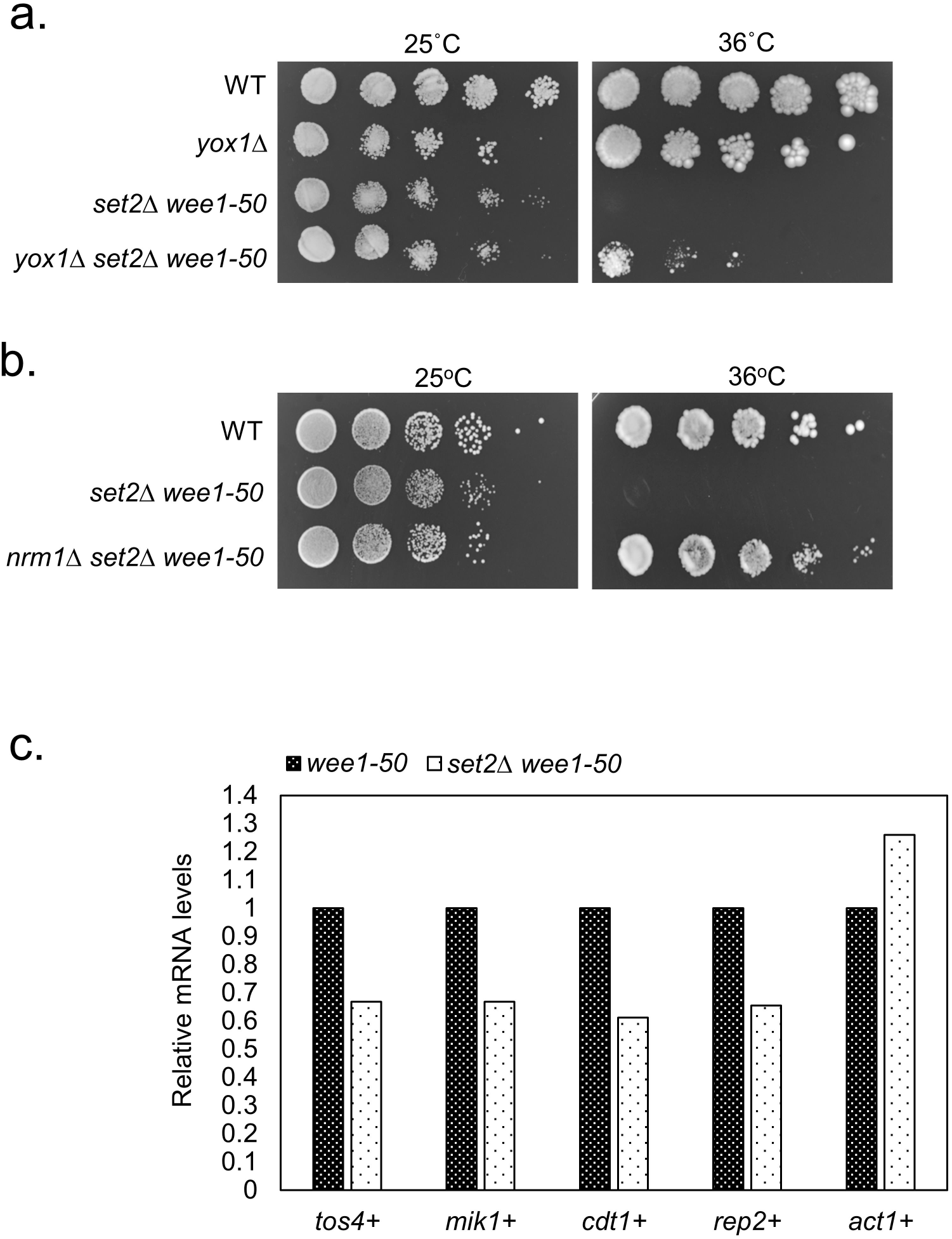
Set2 is required for MBF-dependent gene expression in *wee1-50* cells. (**a**) *yox1Δ* suppresses the synthetic lethality of *set2Δ wee1-50*. Strains were serially diluted and spotted onto YES plates and incubated at 25°C or 36°C for 2-3 days. (**b**) *nrm1Δ* suppresses the synthetic lethality of *set2Δ wee1-50*. Strains were serially diluted and spotted onto YES plates and incubated at 25°C or 36°C for 2-3 days. (**c**) *tos4*^*+*^, *mik1*^*+*^, *cdt1*^*+*^, and *rep2*^*+*^ transcript levels in *set2Δ wee1-50* cells relative to *wee1-50* where the expression level in *wee1-50* cells is 1.0. Data were calculated from two biological repeats. *act1*^*+*^ was shown as an MBF-independent control.

Consistent with observations above (Fig 1a), inactivation of Wee1 also leads to more origin firing in *set2Δ* cells (**Supplementary Fig. 7a**). We also found that partial inactivation of replication licensing factor Cdc18 (*cdc18*^*ts*^ at 34°C) suppressed the synthetic lethality of *set2Δ wee1-50* mutants at the semi-restrictive temperature (**Supplementary Fig. 7b**), suggesting that reducing the number of active replication origins alleviates dNTP depletion in the *set2Δ wee1-50* background. No further synthetic lethality or sickness in *cdc18*^*ts*^ *wee1-50* or *mcm4tdts wee1-50* cells at the semi-restrictive temperature suggesting replication stalling is due to dNTP depletion rather than defects in other steps of DNA replication (**Supplementary Fig. 7c)**. Moreover, we did not observe synthetic sickness between Polε and Wee1 inactivation, indicating that the slow S-phase in *set2Δ* cells is unlikely to be due to the defective polymerase function (**Supplementary Fig. 7d**).

**Figure 7.**
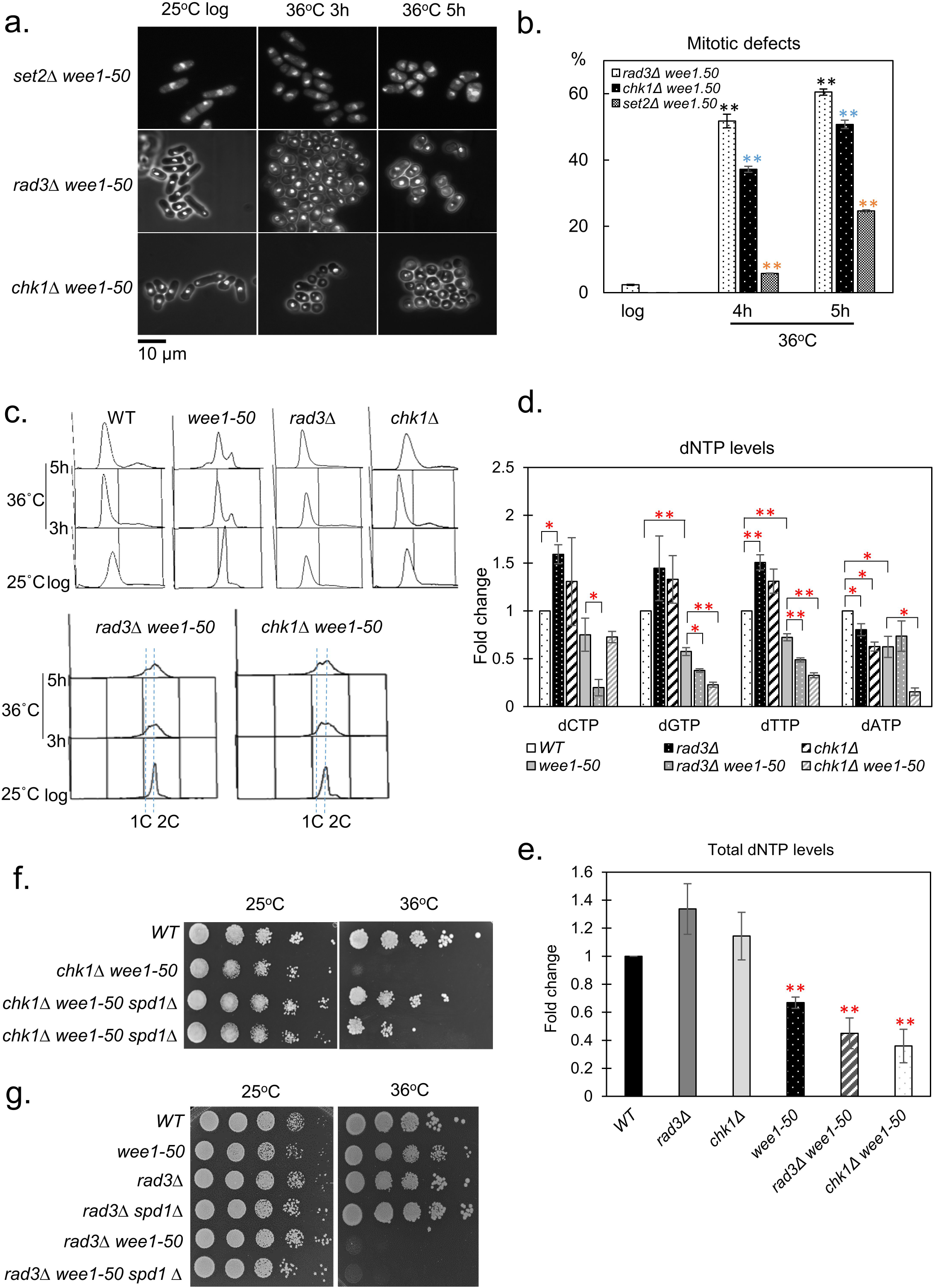
*rad3Δ* and *chk1Δ* are synthetic lethal with *wee1-50* through replication stress. (**a**) *set2Δ wee1-50, rad3Δ wee1-50* or *chk1Δ wee1-50* result in premature entry into mitosis but the ‘’cut’’ phenotype in the set2*Δ wee1-50* mutant is significantly lower at early time points. *set2Δ wee1-50, rad3Δ wee1-50* or *chk1Δ wee1-50* cells were grown to log phase at permissive temperature (25°C), then incubated at 36°C to inactivate Wee1. Samples were fixed with 70% ethanol at indicated times. The fixed cells were stained with DAPI and examined by microscopy analysis. (**b**) Quantitative analysis of cells in (**a**) Means ± standard errors are shown. Black asterisks indicates statistically significant differences between *rad3Δ wee1-50* cells grown at either 36°C or 25°C (** *t* test p<0.01, *t* test p-values for *rad3Δ wee1-50* cells: 4h=0.0017, 5h=0.0003; averages of n ≥ 2 experiments, n ≥100 cells for each data point); blue asterisks indicates statistically significant differences between *chk1Δ wee1-50* cells grown at either 36°C or 25°C (** *t* test p<0.01, *t* test p-values for *chk1Δ wee1-50* cells: 4h=0.0006, 5h=0.0006; averages of n ≥ 2 experiments, n ≥100 cells for each data point); orange asterisks indicates statistically significant differences between *set2Δ wee1-50 cells* grown at 36°C or 25°C (** *t* test p<0.01, *t* test p-values for *set2Δ wee1-50* cells: 4h=0.0002, 5h=0.0078; averages of n ≥ 2 experiments, n ≥100 cells for each data point).(**c**) Wee1 inactivation causes replication stress in *rad3Δ* or *chk1Δ* mutants. Flow cytometric analysis of wild-type, *rad3Δ, chk1Δ, rad3Δ wee1-50* or *chk1Δ wee1-50* cells at 25°C or 36°C at indicated time points. (**d**) dNTP levels were measured in wild-type, *wee1-50, rad3Δ wee1-50* and *chk1Δ wee1-50* strains. These strains were grown to log phase at 25°C following by 5h incubation at 36°C. Samples of cells were collected and re-suspended in 10% TCA for HPLC analysis. Means ± standard errors of three biological repeats are shown. Asterisks (*) indicate statistically significant differences as indicated (p<0.05, *t* test). (e) Total dNTP levels are reduced in *wee1-50, rad3*Δ *wee1-50* or *chk1* Δ *wee1-50* cells compared to wild-type cells. Asterisks (*) indicate statistically significant differences (** *t* test p<0.01, *t* test p-values: *wee1-50*=0.0012, *rad3Δ wee1-50*=0.0075, *chk1Δ wee1-50=* 0.0059).. (**f**) *spd1Δ* suppresses the synthetic lethality of *chk1Δ wee1-50*. Strains were serially diluted and spotted onto YES plates and incubated at indicated temperatures for 2-3 days. (**g**) *spd1Δ* cannot suppress the synthetic lethality of *rad3Δ wee1-50*. A similar experiment was carried out as described in (**f**).

### Disrupting checkpoint-dependent dNTP synthesis with wee1-50 results in replication catastrophe

We and others have identified a role for the DNA damage checkpoint in inducing dNTP synthesis in response to genotoxic stress (Blaikley et al., 2014; Liu et al., 2003; Moss et al., 2010). We therefore compared the effects of inactivatingWee1in*set2Δ*withthoseof*rad3Δ*or*chk1Δ*cells **(supplementary Fig. 8a**, **8b** and **8c)**. We were unable to make the *cds1Δ wee1-50* double mutant as they were lethal at 25°C **(supplementary Fig. 8d)**. We found that *set2****Δ*** *wee1-50* phenocopied the synthetic lethality of *rad3Δ wee1-50, hus1Δ wee1-50* or *chk1Δ wee1-50* mutants **(supplementary Fig. 8a**, **8b** and **8c)**. However, we found that the percentage of cells exhibiting a ‘cut’ phenotype in *set2****Δ*** *wee1-50* cells (20%) was significantly lower compared to *rad3****Δ*** *wee1-50* (60.5%) *or chk1****Δ*** *wee1-50* (50.8%) mutants at 4 or 5h (*p* value <0.05) (Fig. 7a and **7b** and **Supplementary Fig. 9**). We also monitored cell cycle profiles of *rad3****Δ*** *wee1-50* and *chk1****Δ*** *wee1-50* cells following a shift to the restrictive temperature for 5h. Surprisingly, Wee1 inactivation also caused replication stalling in *rad3****Δ*** and *chk1****Δ*** cells (Fig. 7c). In contrast, the *tel1****Δ*** *wee1-50* double mutant did not exhibit synthetic lethality (**Supplementary Fig. 10**), consistent with the fact that *tel1****Δ*** cells exhibited normal S-phase and DNA damage checkpoints (Willis & Rhind, 2009). Together, the above results indicate that disrupting Wee1 causes S-phase arrest in *set2Δ, rad3Δ*, or *chk1Δ* cells, consistent with Wee1 playing an important role in facilitating efficient S-phase progression in fission yeast. We also examined the dNTP levels in the single and double mutants. Unexpectedly, we found that dNTP levels were increased in a *rad3Δ*, or *chk1Δ* cells compared to wild-type cells under unstressed conditions **(Fig.7d)**. This may reflect a lack of DNA damage checkpoint inhibition of the MBF target genes. However, deleting *rad3*^*+*^ or *chk1*^*+*^ in a *wee1-50* background resulted in a significant reduction in dNTP levels compared to wild type or *wee1-50* cells (Fig. 7d). Therefore, these results suggest that Rad3 and Chk1 play an important role in maintaining dNTP levels in the absence of Wee1. Further, we found it was possible to suppress the synthetic lethality following Wee1 inactivation in a *chk1Δ* background by deleting *spd1*^*+*^ in *wee1-50 chk1Δ* cells (Fig. 7e). In contrast, deleting *spd1*^*+*^ did not suppress the synthetic lethality of *rad3Δ wee1-50* double mutants (Fig. 7f), consistent with Rad3 (ATR) playing additional functions in response to replication fork stalling (H. D. Lindsay et al., 1998). These findings together support an essential role for Wee1 in modulating CDK-induced replication stress, and that inactivating Wee1 together with mutations that disrupt dNTP synthesis in response to genotoxic stress results in replication catastrophe.

**Figure 8.**
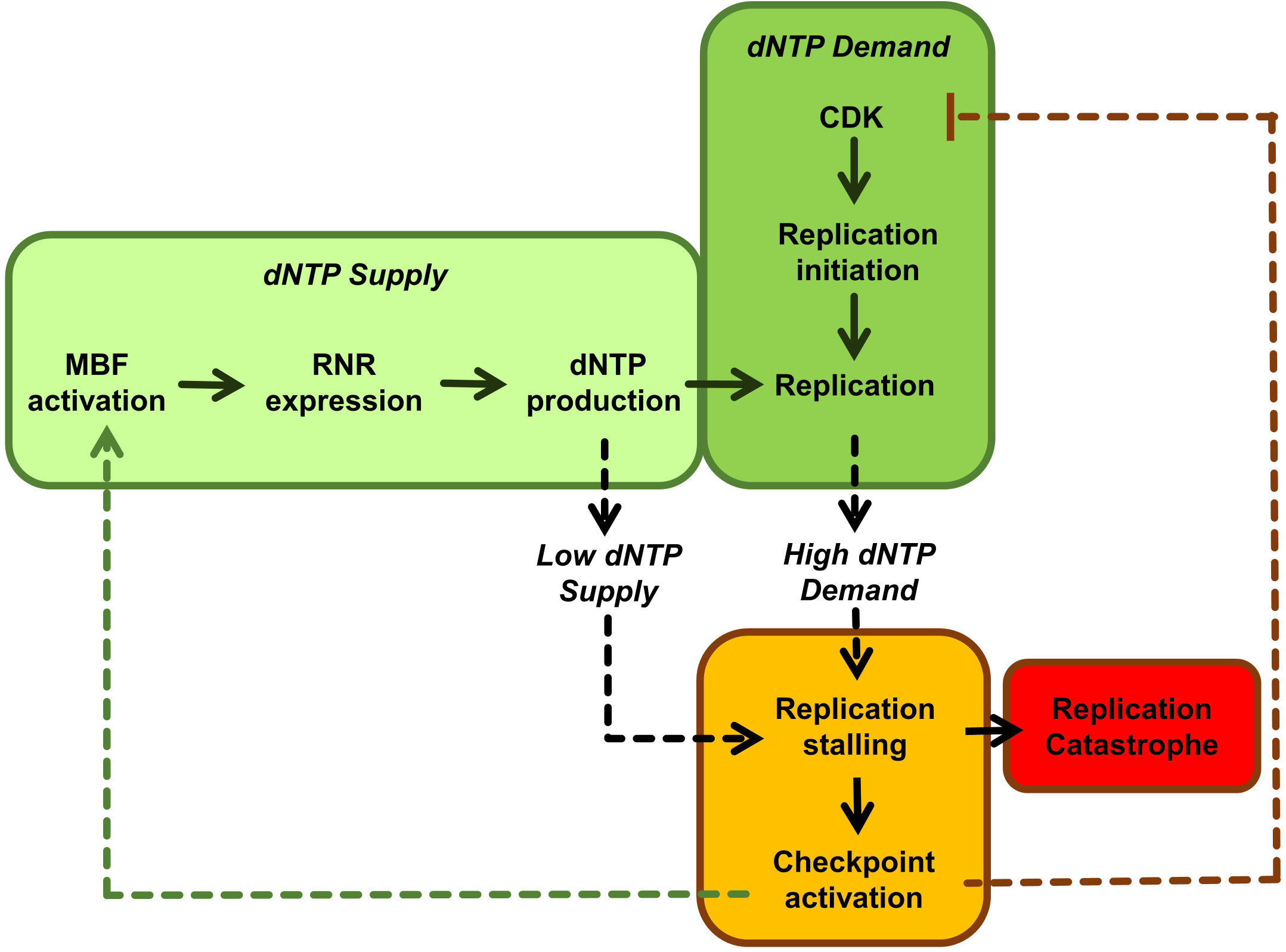
dNTP supply and demand model. Increased CDK activity (resulting from Wee1 inactivation) increases replication origin firing leading to increased dNTP demand (dark green box). This in turn leads to replication stalling and to DNA integrity checkpoint activation (amber box). Checkpoint activation leads to reduced CDK activity and to increased dNTP supply (dotted arrows) through MBF-dependent RNR expression (light green box). Failure to increase dNTP supply (e.g. loss of Set2, Rad3 or Chk1) when dNTP demand is high leads to replication catastrophe (red box). See text for details.

## Discussion

Understanding the mechanisms that can lead to replication stress, and how they can be targeted remains an important goal in cancer research. In this study, we define an evolutionarily conserved role for the CDK regulator Wee1 in suppressing replication stress and dNTP depletion, thereby maintaining genome stability. Further, we demonstrate that dNTP homeostasis defects, resulting from either loss of Set2 or the DNA integrity checkpoint, are synthetic lethal with CDK-induced replication stress, resulting from Wee1 inactivation. Together our results support a ‘dNTP supply and demand’ model, which can be exploited to target replication stress.

Our data indicate that Wee1 inactivation leads to elevated levels of CDK-dependent replication origin firing, resulting in an overall increase in the total number of origins being fired. This in turn leads to dNTP depletion, replication stress, Mus81-dependent DNA damage and subsequent genome instability. We found that the replication stress associated with Wee1 inactivation alone, or in combination with *set2Δ,* resulted in Mus81-dependent Rad52-GFP foci formation. Such Mus81 activity is likely to have been triggered by stalled replication forks, which present as substrates for this structure-specific endonuclease (Osman, Dixon, Doe, & Whitby, 2003). Mus81 is carefully regulated during S phase and G2 to prevent it inappropriately cleaving stalled forks (Froget, Blaisonneau, Lambert, & Baldacci, 2008; Kai, Boddy, Russell, & Wang, 2005). Elevated CDK activity has been shown to promote Mus81 activation through phosphorylation of Eme1 (Dehé et al., 2013) and we hypothesise that elevated levels of CDK in S phase of *wee1-50* cells contributes to the Mus81-dependent DNA damage. We also observed that Wee1 inactivation resulted in robust induction of Cdc22, the catalytic subunit of RNR, thus promoting dNTP synthesis. These findings are consistent with a major role for Wee1 in regulating CDK activity in S-phase in fission yeast (Anda, Rothe, Boye, & Grallert, 2016) and support an evolutionarily conserved role for of WEE1 in regulating dNTP usage and preventing DNA damage through regulating origin firing (Beck et al., 2012). Our findings further demonstrate that Wee1 inactivation has significant consequences for genome stability.

We find Wee1 inactivation together with loss of Set2-dependent histone H3K36 tri-methylation results in synthetic lethality. Our data support a key role for Set2-dependent H3K36me3 in facilitating MBF-dependent Cdc22 transcription and thus promoting dNTP synthesis in response to genotoxic stress (Pai et al., 2017). Further, Set2 dependent dNTP synthesis becomes essential following Wee1 inactivation and CDK-induced dNTP depletion. In support of this, we find loss of Set2 reduces MBF-dependent Cdc22 expression, the catalytic subunit of RNR, and leads to dNTP pool depletion in response to genotoxic stress. Simultaneous loss of Wee1 and Set2 leads to critically low dNTP pools and a failure to induce Cdc22 expression and to subsequently replenish dNTP levels following Wee1 inactivation. This in turn leads to cell death through replicative arrest and mitotic catastrophe. Consistent with this, we find that *set2Δ wee1-50* synthetic lethality is associated with S-phase arrest; Cdc22 expression is significantly reduced in the double mutant compared to wild-type; dNTP levels are significantly reduced in the double mutant compared to *wee1-*50, and the double mutant is acutely sensitive to HU at the permissive temperature. Accordingly, the synthetic lethality can be suppressed through increasing dNTP synthesis by depleting Spd1, by increasing MBF-dependent Cdc22 expression, or by compromising replication origin licensing. The fact that *mus81*^+^ deletion did not robustly suppress the *set2Δ wee1-50* synthetic lethality is consistent with Mus81 cleavage of collapsed forks being a downstream secondary consequence of dNTP depletion, which are the primary cause of cell death.

We further define a more general role for dNTP synthesis in maintaining viability in response to Wee1 inactivation. Wee1 inactivation has been previously found to be synthetic lethal with loss of Rad3 (ATR) or Chk1 in both yeast and humans (al-Khodairy & Carr, 1992; Enoch et al., 1992; Srivas et al., 2016). Synthetic lethality between Wee1 and checkpoint deficient mutations has been proposed to be a consequence of mitotic catastrophe. However, our results demonstrate that, while mitotic catastrophe is observed in *rad3Δ* or *chk1Δ* checkpoint mutants following Wee1 inactivation, these cells undergo prior replication arrest resulting from an insufficient dNTP supply. Importantly, the Rad3 (ATR)-dependent checkpoint pathway is required to induce dNTP synthesis following replication stress and DNA damage. DNA damage checkpoint activation leads to Cul4-Ddb1^Cdt2^ dependent degradation of Spd1, a negative regulator of RNR to promote dNTP synthesis (Moss et al., 2010). The replication checkpoint also promotes MBF-dependent transcription of Cdc22, the catalytic subunit of RNR through Cds1-dependent phosphorylation of Yox1, which blocks the binding of this negative regulator to MBF in response to replication stress (Ivanova, Gomez-Escoda, Hidalgo, & Ayte, 2011). In this respect, Set2 and the DNA integrity checkpoint function analogously to facilitate dNTP synthesis in response to both DNA damage and replication stress in fission yeast. Accordingly, we show that elevating dNTP levels by deletion of *spd1*^*+*^ suppressed the synthetic lethality of both the *set2Δ wee1-50* and *chk1Δ wee1-50* mutants. That *spd1*^*+*^ deletion could not suppress the synthetic lethality of *rad3Δ wee1-50* mutant likely reflects the fact that Rad3 (ATR) performs additional roles in replication fork restart. Together, these findings support a ‘dNTP supply and demand’ model in which Set2 and DNA integrity checkpoint dependent dNTP synthesis becomes essential following elevated CDK-induced origin firing and dNTP depletion, thereby preventing replication catastrophe. This model explains how Wee1 inactivation results in synthetic lethality with loss of Set2; sheds new light on the synthetic lethal relationship between loss of ATR, Chk1 and Wee1 inactivation; and further predicts that other mutations that disrupt dNTP synthesis in response to replication stress will also be synthetic lethal with Wee1 inactivation (Figure 8).

Our findings indicate that the S-M cell cycle checkpoint is intact in *set2Δ* cells (Pai et al., 2017), where an elongated phenotype being observed in response to hydroxyurea or bleomycin. Moreover, in contrast to *rad3Δ* or *chk1Δ*, Wee1 inactivation did not lead to rapid mitotic catastrophe in *set2Δ* cells. While *set2Δ wee1-50* cells underwent mitotic catastrophe at later time points, this may reflect a role for Set2 in promoting MBF-dependent transcription of *mik1*^*+*^, encoding Mik1 kinase, which negatively regulates Cdc2 and leads to mitotic catastrophe when deleted in a *wee1-50* background (Christensen, Bentley, Martinho, Nielsen, & Carr, 2000; Dutta et al., 2008; Dutta & Rhind, 2009; Lee, Enoch, & Piwnica-Worms, 1994; Lundgren et al., 1991; Ng, Anderson, White, & McInerny, 2001).

Based on findings described here, it was demonstrated that H3K36me3-deficient human cancers are synthetic lethal with the WEE1 inhibitor AZD1775 as a result of dNTP starvation (Pfister et al., 2015). These findings are of clinical relevance as despite the frequent loss of histone H3K36me3 in multiple cancer types and its association with poor patient outcome, there is no therapy targeting H3K36me3-deficient cancer types (Forbes et al., 2015; Lawrence et al., 2014; Li et al., 2016). Moreover, our data suggests that inhibitors of ATR and CHK1 may have differential effects in cancer therapy. As inhibitors to WEE1, ATR and CHK1 are already in clinical trials (http://www.clinicaltrials.gov), we anticipate that our findings described here will provide important mechanistic insights into the targeting of cancers exhibiting replication stress.

## MATERIALS AND METHODS

### Yeast strains, media and genetic methods

The strains used in this study are listed in Supplementary Table I. Standard media and growth conditions were used. Cultures were grown in rich media (YE6S) or Edinburgh minimal media (EMM) at 32?°C with shaking, unless otherwise stated. Nitrogen starvation was carried out using EMM lacking NH_4_Cl.

### Serial dilution assay

A dilution series for the indicated mutant cells was spotted onto YES plates. Plates were incubated at 25?°C, 32?°C or 36?°C for 2-3 days, as indicated, before analysis.

### Survival analysis

Exponential cultures were obtained in liquid YE6S medium inoculated with a single colony picked from a freshly streaked (YE6S) stock plate and grown overnight at 25°C with vigorous shaking. Exponential cells were resuspended in YE6S at a density of 2 × 10^7^ cells ml^-1^. Serial dilutions were made and 500 cells were plated on YE6S plates at the restrictive temperature of 36°C, as well as a control plate incubated at 25°C. Plates were incubated for 2-3 days and colonies were then scored.

### Analysis of replication origin firing

The polymerase usage sequence (Pu-seq) technique was performed as previously described (Daigaku et al., 2015). Briefly, DNA was extracted from cells grown to log phase either on 18°C or on 34°C as indicated. For ‘wt’ datasets two strains were used, both strains containing *rnh201* deletion together with either polymerase δ (*cdc6-L591G*) or polymerase ε (*cdc20-M630F*) mutations. These strains incorporate more rNTPs on the strands synthetized by the mutant polymerase. These sites can be mapped by Pu-seq. For the *wee1-50*, and *wee1-50 set2Δ* datasets the two strains also contained these mutations along with *rnh201* and *cdc6-L591G* or *cdc20-M630F*. The isolated DNA was then subjected to alkali treatment (0.3 M NaOH, 2h 55°C) which digested the DNA at the positions of rNTP incorporation and also separated the double strands. The resulting ssDNA fragments were size selected on agarose gel (fragments between 300-500bp were isolated). These fragments were then used for creating strand specific next generation sequencing libraries and sequenced on a Next-seq Illumina platform resulting in ∼10M reads form each strains. The Pu-Seq data has been uploaded to Gene Expression Omnibus (GEO); accession number GSE113747. Reads were aligned to the *Schizosaccharomyces pombe* reference sequence (http://www.pombase.org/downloads/genome-datasets), the reads were mapped using bowtie2 and the data was analyzed and origin positions and efficiencies were determined using the tools published and described in detail in Daigaku et al., 2015 with default variables except for the ‘percentile threshold for origins’ option was set to 0.2 = 20th percentile. Efficient origins were determined as origins with higher than 50% efficiency and inefficient origins had less than 25% efficiency.

### Mini-chromosome instability assay

The mini-chromosome loss assay was carried out as previously described (Allshire, Nimmo, Ekwall, Javerzat, & Cranston, 1995; Moss et al., 2010). Briefly, 500–1000 cells from individual Ade^+^ colonies were plated on EMM plates containing low adenine (5 mg/L), incubated at 25°C, 30°C or 36°C for 3 days and were stored for 48h at 4°C before being scored for the presence of sectored colonies. The number of mini-chromosome loss events per division was determined as the number of Ade^-^ sectored colonies divided by the sum of white and sectored colonies. The experiment was performed in triplicate.

### The *CanR* mutation assay

To analyse mutation rates, a Luria-Delbruck fluctuation analysis was performed (Luria & Delbruck, 1943). Briefly, 1 mL cultures of wild-type or *wee1-50* cells were grown in YES medium to saturation in 12-well plates at 25°C. 100 µL of each culture was spotted onto PMG (-arg, -his) plates containing 100 µg/mL canavanine and incubated at 32°C for 10-12 days. Colony numbers were scored and mutation rates in culture were analysed using the FALCOR tool (http://www.keshavsingh.org/protocols/FALCOR.html) (Hall, Ma, Liang, & Singh, 2009). For each strain, colony data were collected from at least 30 independent cultures. Averages, standard deviations and error bars were calculated for three independent experiments.

### Microscopy analysis

Asynchronous cell cultures were treated with 10 mM hydroxyurea (HU) at the indicated temperature before being fixed in methanol. Samples were rehydrated and stained with 4′,6-diamidino-2-phenylindole (DAPI) before examination using Zeiss Axioplan 2ie microscope, Hamamatsu Orca ER camera and micromanager software. For visualization of Rad22-GFP foci, cells were incubated at 25°C or 32°C for 5 hours before being fixed and visualized as above.

### Protein analysis

Protein extracts were made by TCA extraction and analyzed by Western blotting as described previously (Pai et al., 2014). TAP-tagged proteins were detected with peroxidase–anti-peroxidase–soluble complex (P1291, Sigma). Cdc22-GFP was detected using antibody anti-GFP (11814460001, Roche), and α-tubulin was detected with antibody T5168 (Sigma).

### dNTP analysis

10^8^ cells were collected and washed with 2 % glucose. Cell pellets were then lysed with 50 µl 10 % TCA and stored at −80°C before HPLC analysis. On thawing, cell extracts were spun and the supernatant diluted five-fold with water. Samples were then neutralised and analysed by HPLC as described by Moss et al using a Waters e2695 autosampler. All peak areas were measured at 258 nm (Moss et al., 2010).

### Microarray analysis

Microarray analysis was performed as previously described (Pai et al., 2014; Rallis, Codlin, & Bahler, 2013). Experiments were conducted in duplicate with a dye swap. RNAs from two independent biological replicates have been utilised for cDNA production. Figure 6c shows average expression ratios from the two repeats. Original data are deposited in ArrayExpress; accession number E-MTAB-6795. In brief, Alexa 555- or 647-labeled cDNA was produced from the RNA, using a Superscript direct cDNA labelling system (Invitrogen) and Alexa 555 and 647 dUTP mix. cDNAs were then purified using an Invitrogen PureLink PCR Purification system and hybridized to the array using a Gene Expression Hybridization kit (Agilent). The arrays are Agilent custom-designed containing 60-mer oligonucleotides synthesized *in situ* containing 15,000 probes. Following hybridization for at least 17 hours, the arrays were washed using a Gene Expression Wash Buffer kit (Agilent) and scanned in an Agilent Array Scanner. Signals were extracted using GenePix software.

## ACKNOWLEDGEMENTS

C.C.P, K.F.H, S.C.D, A.K, A.M.C. S.E.K, C.R, L.K.F, R.D and N.D.L performed and analysed the experiments. C.C.P and T.C.H wrote the manuscript with input from all authors. Experiments in Fig. 1 were performed by C.C.P, A.K and A.M.C; in Fig. 2 were performed by C.C.P and S.E.K; in Fig. 3 were performed by C.C.P, K.F.H, and C.J.S; in Fig. 4 were performed by C.C.P and S.C.D.; in Fig. 5 were performed by C.C.P, S.C.D and L.K.F; in Fig. 6 were performed by C.C.P, C.R and J.B; in Fig. 7 were performed by C.C.P, N.D.L and L.K.F.

## SUPPLEMENTARY FIGURE LEGENDS

**Supplementary Figure 1** *wee1-50* cells are sensitive to HU. (a) Equal amount of cells were streaked on YES or 10 mM HU and incubated at 32°C. (b) dATP levels in *wee1-50* cells were significantly lower than wild-type cells. dATP levels were normalised with ATP levels. The asterisk (*) represents significant difference compared with wild type and *wee1-50* (p < 0.05, t test).

**Supplementary Figure 2** Expression of different levels of human histone demethylase in *S. pombe*. (a) 10-fold serial dilutions of wild type expressing empty vector, hJMJD2A using three different *nmt* promoters were spotted onto EMM minus leucine without thiamine at 36°C. (b) 10-fold serial dilutions of wild type expressing empty vector, hFBXL11 using three different *nmt* promoters were spotted onto EMM minus leucine without thiamine at 36°C. (c) Western blot analysis of overexpression of FLAG-hJMJD2A levels in wild-type cells under the control of *pREP3X, pREP41X* and *pREP81X* plasmids. Antitubulin as a loading control. (d) Western blot analysis of overexpression of FLAG-hFBXL11 levels in wild-type cells under the control of *pREP3X, pREP41X* or *pREP81X* plasmid. Anti-tubulin is shown as a loading control. (e) Western blot analysis of hJMJD2A levels in wild-type or *wee1-50* cells containing *pREP41x-Flag-FBXL11* plasmids. α-tubulin is shown as a loading control. (f) Western blotting analysis of hFBXL11 levels in wild-type or *wee1-50* cells containing *pREP41x-Flag-FBXL11* plasmids. α-tubulin is shown as a loading control.

**Supplementary Figure 3**Loss of Clr4 is not essential for the viability of *wee1-50* cells. Serial dilution of a wild-type, *wee1-50, clr4Δ* and *clr4Δ wee1-50* strains were spotted onto YES medium and incubated at indicated temperatures for 2-3 days.

**Supplementary Figure 4** Loss of Cdc2 function rescues the synthetic lethality of *set2Δ wee1-50* cells. (a) Serial dilution of a wild-type, *cdc2-as, set2Δ wee1-50* and *set2Δ wee1-50 cdc2-as* strains were spotted onto YES medium and incubated at indicated temperatures for 2-3 days. (b) Quantification of the cell viability of indicated strains at 36°C.

**Supplementary Figure 5** *set2Δ wee1-50* cells exhibit severe mitotic catastrophe at later time points following Wee1 inactivation. (a) *set2Δ wee1-50* cells were grown to log phase at the permissive temperature then transferred to the restrictive temperature for 24h. Samples of cells were collected at indicated time points and stained with DAPI for microscopy analysis. (b) Quantification of mitotic defects in (a).

**Supplementary Figure 6** *set2Δ wee1-50* cells exhibit proficient checkpoint activation in response to HU. Cells were grown asynchronously in YES medium and transferred to YES in the present of 10mM HU for 24h at 25°C. Samples were taken at the indicated time points and fixed with methanol/acetone; subsequently the fixed cells were examined by microscopy analysis.

**Supplementary Figure 7** Inhibition of origin firing suppresses the synthetic lethality of *set2Δ wee1-50* cells. (a) Set2 and Wee1 suppresses inefficient origin firing. The genome-wide plot of origin usage in vegetative wild-type, *set2Δ, wee1-50* or *set2Δ wee1-50* cells at 34°C. Origin efficiencies were calculated from Pu-seq data. (b) Inactivation of Cdc18 at the semi-restrictive temperature suppresses the synthetic lethality of *set2Δ wee1-50* cells (c) Inhibition of replication factors Cdc18 or Mcm4 does not cause severe viability loss with Wee1 inhibition in *S. pombe. cdc18*^*ts*^ or *mcm4-tdts* is not synthetic lethal with *wee1-50*. (d) *pol?*^*ts*^ is not synthetic lethal with *wee1-50*.

**Supplementary Figure 8** Checkpoint mutant is synthetic lethal with *wee1-50* mutant. (a) *rad3Δ* is synthetic lethal with *wee1-50*. Serial dilution of a wild-type, *rad3Δ, wee1-50* and *rad3Δ wee1-50* strains were spotted onto YES medium and incubated at indicated temperatures for 2-3 days. (b) *hus1Δ* is synthetic lethal with *wee1-50*. Similar experiments were carried as described in (a). (c) *chk1Δ* is synthetic lethal with *wee1-50*. Similar experiments were carried as described in (a). (d) *cds1Δ* is synthetic lethal with *wee1-50* at 25°C.

**Supplementary Figure 9** Wee1 inhibition causes pre-mature entry into mitosis in checkpoint mutants. Percentage of septated cells flowing Wee1 inhibition in *rad3Δ, chk1Δ* or *set2Δ* cells.

**Supplementary Figure 10** *tel1Δ* is not synthetic lethal with *wee1-50*. Serial dilution of a wild-type, *tel1Δ, wee1-50* and *tel1Δ wee1-50* strains were spotted onto YES medium and incubated at 25°C or 36°C for 2-3 days.

